# A Post-hurricane Quantitative Assessment of the Red-bellied Racer (*Alsophis rufiventris*) on Saba and Comparison with St. Eustatius

**DOI:** 10.1101/2021.11.15.468688

**Authors:** Hannah Madden, Lara Mielke

## Abstract

We estimated occupancy, abundance (lambda), detection probability, density/ha and abundance of a regionally endemic snake in the Colubrid family on the Dutch Caribbean island of Saba in 2021, four years after hurricanes Irma and Maria impacted the island. Line transect surveys were conducted at 74 sites covering 6.7 ha. The proportion of sites occupied was estimated at 0.74 (min 0.48, max 0.90), with occupancy varying between vegetation types and across elevational gradients. Similarly, lambda was estimated at 1.61 (min 0.7, max 3.7) but varied between vegetation types and elevational gradients. Detection probability was estimated at 0.15 (min 0.10, max 0.21). Using Distance sampling, we estimated 10.9 (min 7.3, max 16.2) racers/ha, with a total population estimate of 4,917 (min 2,577, max 6,362) across the entire study region (438.6 ha.) Based on anecdotal observations from Saban residents and prior literature describing the pre-hurricane population as “abundant” (at least 2.0 racers/hour), we posit that the population experienced a hurricane-induced decline but may have since recovered, though not to previous levels (1.28 racers/hour). Nevertheless, our results suggest that racer densities on Saba are currently higher than those on St. Eustatius. Despite this, given the species’ extremely limited extant range and the presence of invasive species on both islands, prevention of local extirpation should be a high conservation priority.

## INTRODUCTION

Reptile populations in the tropics are influenced by island size (Ricklefs and Lovette 1999), elevation (Fauth et al., 1989; Ricklefs and Lovette, 1999), geography (Moser et al., 2018), topography (Ricklefs and Lovette 1999) and vegetation structure (Herrera-Montes and Brokaw 2010). Furthermore, reptiles face a multitude of threats, including but not limited to predation by invasive species (Gibbons et al., 2000), habitat disturbance/loss (Mayani-Parás et al., 2019), human disturbance/persecution (Alves et al., 2012; Atwood et al., 2020), and climate change (Urban, 2015). In the Caribbean, hurricanes in particular are predicted to increase in frequency and severity by the end of the 21^st^ century (Webster et al., 2005; Bender et al., 2010). The impacts of increased hurricane activity and severity on reptile populations are as yet unknown, and as stated by Powell (2004), “we can only guess at changes resulting from …. the impact of hurricanes on populations occupying increasingly smaller parcels of suitable habitats”.

The Caribbean Region is a biodiversity hotspot due to its high levels of species endemism (Mittermeier et al., 2004); unfortunately, their limited geographic ranges makes species more vulnerable to extinction from local impacts (e.g., habitat loss, invasive species; Catford et al., 2012). The Caribbean Netherlands volcanic island of Saba (13 km^2^) is dominated by a single dormant volcano, Mount Scenery, which reaches a maximum elevation of 870m above sea level. The climate at lower elevations is described as tropical savannah, whereas the upper slopes of Mount Scenery are characterized as a tropical rainforest climate (de Freitas et al., 2016). The island’s complex topography has created a variety of vegetation types that are influenced by changes in wind, sun, and rain exposure (de Freitas et al., 2016). Vegetation on the western side of Saba is predominantly deciduous forest, the mid-latitude southern side of the mountain is tropical rainforest, and the northeastern side is herbaceous rangeland (Stoffers, 1956). However, Saba’s natural vegetation has changed extensively since it was first described by Stoffers (1956). These changes are thought to be attributed to hurricane impacts, natural succession due to reduced agricultural activity, and invasive species (de Freitas et al., 2016). In September 2017 two category 5 hurricanes (Irma and Maria) struck the Windward Islands, causing extensive damage to forests on Saba. Tree stems were defoliated, branches were lost, and average mortality was estimated at 18% (Eppinga and Pucko, 2018). Natural systems are thought to require decades to recover from the impacts of a category 5 hurricane (Lugo, 2000). Studies on the impacts of hurricanes on wildlife populations in the tropics are generally lacking (but see Rivera-Milán et al., 2021; Madden et al., 2021; van den Burg et al., 2021).

West Indian racers in the family Colubridae occur throughout the Lesser Antilles but are in rapid decline (Henderson, 1992, 2004; Powell and Henderson, 2005). Racers (*Alsophis* spp.) are predominantly ground-dwelling, diurnal (but see Madden, 2020 and Questel, 2021), oviparous species that are susceptible to invasive predators such as black rats (*Rattus rattus* Linnaeus, 1758), domestic cats (*Felis catus* Linnaeus, 1758) and mongooses (*Herpestes javanicus* Geoffroy Saint-Hilaire, 1818; Daltry et al., 1997; Powell, 2006; Debrot et al., 2014). Racers are especially vulnerable as they can invoke (irrational) fear among humans, leading to persecution, and due to habitat degradation (Powell, 2006). For a brief overview of *Alsophis* spp. in the Caribbean and a full description of the current status of *Alsophis rufiventris* (Duméril, Bibron and Duméril, 1854) on St. Eustatius, see Madden et al. (2021).

Quantitative assessments of racers are generally lacking (but see Daltry et al., 2017 and Hileman et al., 2017), as highlighted by Madden et al. (2021), and to our knowledge, only one study has quantified the effects of hurricanes on *A. rufiventris* on St. Eustatius (Madden et al., 2021). Specifically, the authors documented a significant decline in encounter rates of *A. rufiventris* on St. Eustatius (16.0 snakes/hr in 2011 compared to 0.34/hr in 2018 and 0.41/hr in 2019) as a result of hurricanes Irma and Maria. While no systematic pre-hurricane data exist from Saba, Sajdak and Henderson (1991) stated that “population density at higher altitudes (>500 m) on Saba was the greatest of any racer population we have seen”. Similarly, Daltry et al. (1997) described the racer as “more abundant on Saba than St. Eustatius (e.g. one racer was seen every thirty minutes whilst walking along paths through the Saban rainforest, whereas sightings in St Eustatius occurred much less frequently”). However, the authors stressed the importance of recognizing natural seasonal variations in apparent abundance of the species. Mating is thought to commence in March, whereby males would be more active and thus more readily detected. Racer populations are also thought to fluctuate with rainfall patterns (Daltry et al., 1997). The species was previously described as “thriving” on Saba (and St. Eustatius), where it was common even in urban areas (Henderson, 2004; Maley et al., 2006). Thus, if pre-hurricane racer populations were indeed higher on Saba than St. Eustatius, we might assume that Daltry et al.’s (1997) count of 2/hr was a significant under-estimation. This study aims to rectify the lack of quantitative assessments of the racer on Saba by providing occupancy, lambda, detection probability, density and abundance estimates of the population within a specific stratum.

## METHODS

### Study Area

Our study took place on Saba (17°37’N, 63°14’ W), Caribbean Netherlands, a small (13 km^2^) island with a human population of approximately 1,933 (CBS 2021) located approximately 30.6 km north-east of St. Eustatius (Fig. 1). Temperature typically ranges from 24 to 31°C. Average annual rainfall is 760.5 mm, with the upper slopes of Mount Scenery receiving >2,000 mm per year (Veenenbos, 1955; de Freitas et al., 2016).

**Figure 1.**
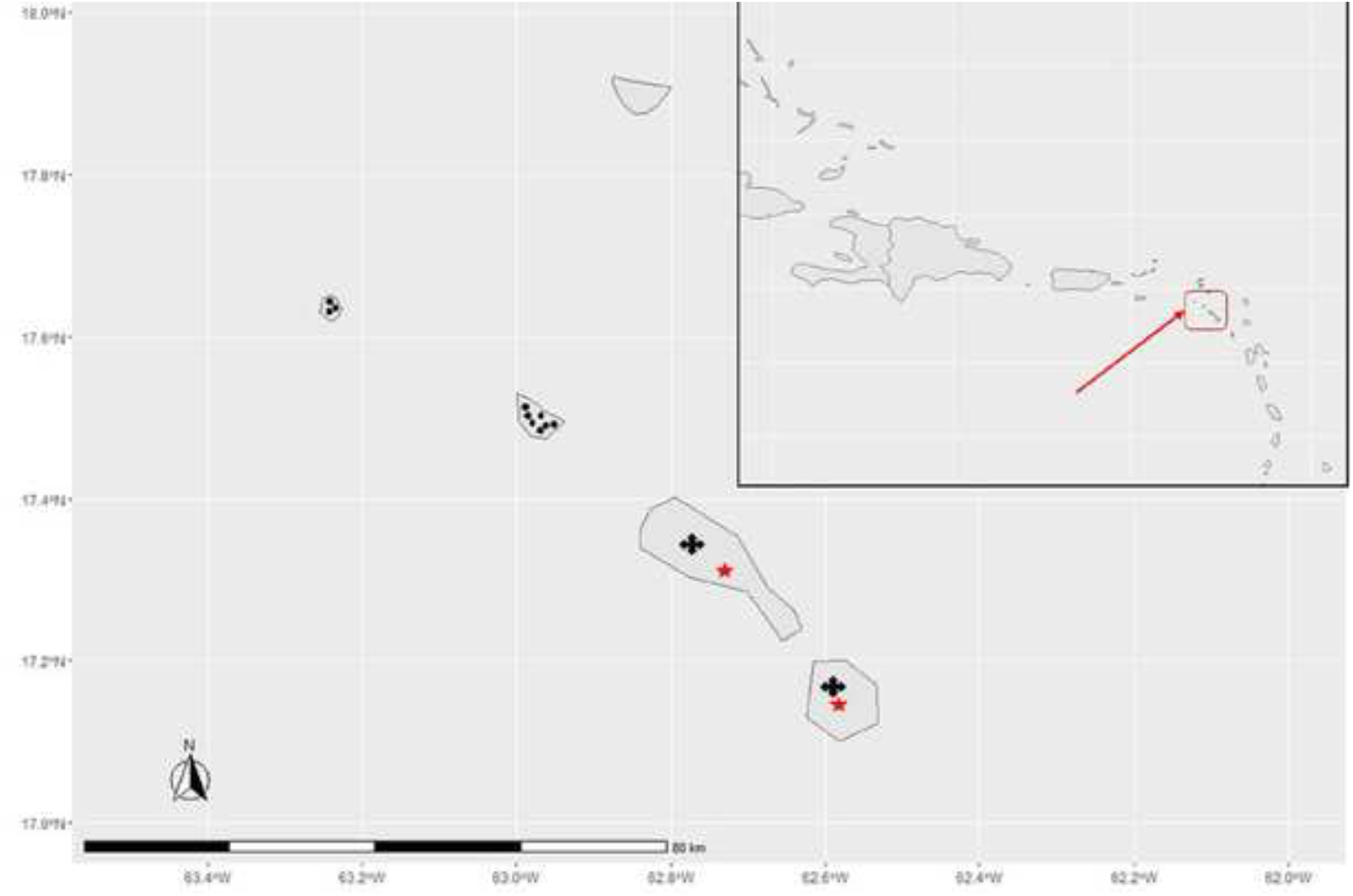
Distribution of *Alsophis rufiventris* on the St. Christopher and Saba banks. Dots indicate locality records, X’s mark extirpated populations on St. Kitts and Nevis, and red stars mark fossil localities (adapted from Maley et al., 2006). Red arrow indicates the location of Saba.

### Post-hurricane Surveys

Between July and September 2021 we surveyed racers in six vegetation types on Saba (total survey area 6.7 ha). To maximize detection and minimize disturbance, all surveys were conducted along existing hiking trails. Using the existing network of trails allowed us to survey racers in an otherwise steep topographical environment with, at times, rocks or dense vegetation. We conducted surveys along 100 m line transects by looking and listening for *A. rufiventris* while walking at a very slow pace. All transect surveys were conducted between 0800 and 1800 hrs. Upon detection, the perpendicular distance of the snake from the center of the transect was measured with a tape measure. Elevation (m) was measured with a handheld GPS.

#### Habitat descriptions

We conducted fieldwork in six habitat types based on vegetation descriptions by de Freitas et al. (2016; Fig. 2): M1 (*Heliconia-Charianthus* Mountains; the highest parts of Mount Scenery, 770-870m), M2 (*Philodendron-Marcgravia* Mountains; the lower slopes of Mount Scenery, 360-860m), M3 (*Philodendron-Inga* Mountains; hilltops around Mount Scenery, 110-810m), M4 (*Swietenia* Mountains; lower western slope of Mount Scenery, 2-270m), M6 (*Coccoloba-Inga* Mountains; south-west side of Saba, 3-550m), and M7 (*Aristida-Bothriochloa* Mountains, 0-500m).

**Figure 2.**
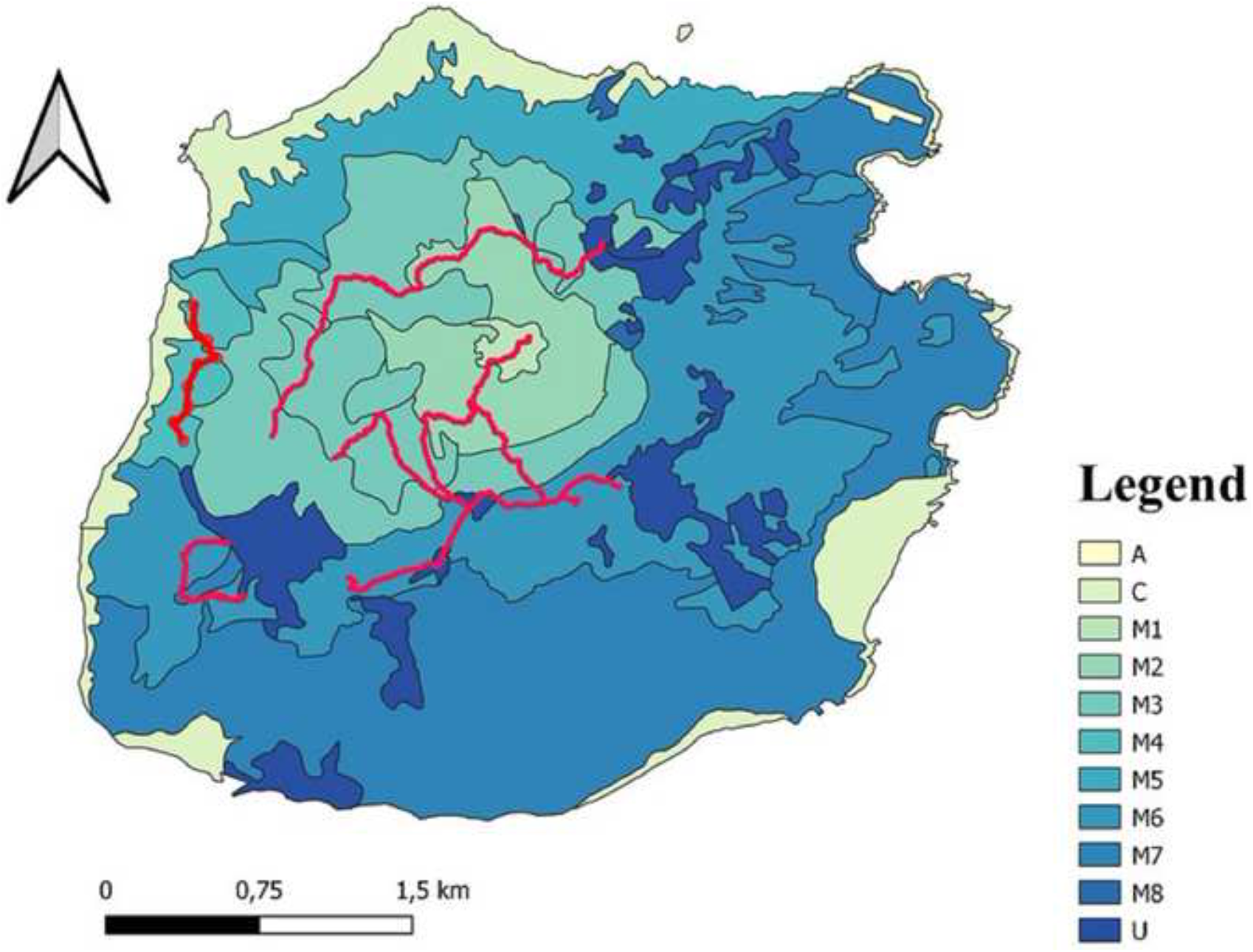
Map of Saba showing vegetation types (M1 – M8) and line transects (red lines). The remaining codes stand for A = Airport, C= Cliffs, U = Urbanized areas (adapted from de Freitas et al., 2016).

### STATISTICAL ANALYSES

We used a likelihood-based, single-season occupancy model (MacKenzie et al., 2017) in the package ‘wiqid’ (Meredith, 2020). We followed the methods described by Madden et al. (2021) to obtain estimates of site occupancy 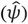 in relation to habitat and elevation, average number of individuals per sample, lambda 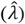, and sampling detection probability 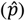. We also used the package ‘Rdistance’ (McDonald et al., 2019) to obtain density (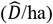 and abundance 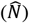 estimates of racers on Saba and St. Eustatius (in the latter case by pooling two years’ of data, which provided annual estimates) across both survey regions (458.6 ha on Saba and 540 ha on St. Eustatius). Due to the very small number of detections on each island it was not possible to separate the data into smaller strata (e.g., vegetation types). Distance sampling is based on estimation of a detection function, 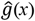, which for line transects decreases with distance and is required to estimate detection probability of individuals or clusters of individuals in the survey region (Buckland et al. 2001, 2015). Following exploratory data analysis (Buckland et al. 2001, 2015), we right truncated the distance data (*w* = 2.5 m) and estimated detection probability of available individuals and clusters. We assessed the fit of uniform, half-normal, and hazard-rate detection models with goodness of fit tests (e.g., Kolmogorov-Smirnov test D_*n*_, P < 0.05); and we ranked the resulting models using Akaike Information Criterion (AIC) for stepwise model selection (Buckland et al., 2001, 2015, Marques et al. 2007). We used nonparametric bootstrapping (500 simulations) to estimate detection probability standard errors (Efron and Tibshirani, 1993, Buckland et al., 2001, 2015). All analyses were performed in the R environment version 4.0.3.

## RESULTS

### Post-hurricane Survey Population Size

We conducted surveys between 4 July and 28 September 2021, four years after Hurricanes Irma and Maria, surveying an area of 6.7 ha in six vegetation types. Average transect length was 100.55 ± SD 5.72 m. Line transects were repeated a minimum of four and a maximum of 12 times at different sites. Average survey time per transect was 5.50 ± 2.73 minutes. We detected a total of 56 racers during surveys.

Occupancy modeling revealed 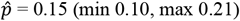, which increased to 0.17 (min 0.08, max 0.31) when a racer was previously detected. Average 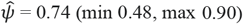, with estimates varying between habitats (Fig. 3) and increasing in line with elevation (Fig. 4). Our 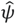 estimate is similar to estimates obtained from St. Eustatius (0.73 in 2018, 0.65 in 2019; Madden et al. 2021), however 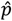 is higher than on St. Eustatius (0.05 in 2018 and 0.07 in 2019; Madden et al. 2021). Occupancy varied across the six habitats sampled, being highest in M1 (peak of Mount Scenery) and lowest in M7 (*Aristida-Bothriochloa* Mountains), though we note the wide 95% confidence intervals for each habitat type, some of which encompass each other. Average 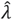 was 1.61 (min 0.70, max 3.67), with estimates varying between habitats (some with very wide and overlapping CIs; Fig. 5) and increasing in line with elevation (Fig. 6), although 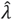 declines steeply above 800m. Estimated encounter rate was 1.28/hr. Mean 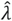 was similar to estimates from St. Eustatius (1.82 in 2018 and 1.60 in 2019). Our raw encounter rate of 1.28/hr is higher than estimates from St. Eustatius (0.34 and 0.41/hr) but lower than the pre-hurricane estimate of 16.0 snakes/hr (Madden et al., 2021), as well as Daltry et al.’s (1997) estimate of 2/hr from Saba. Similarly, our density estimate from Saba of 10.9/ha is higher than those from St. Eustatius (9.9/ha in 2018 and 7.3 in 2019).

**Figure 3.**
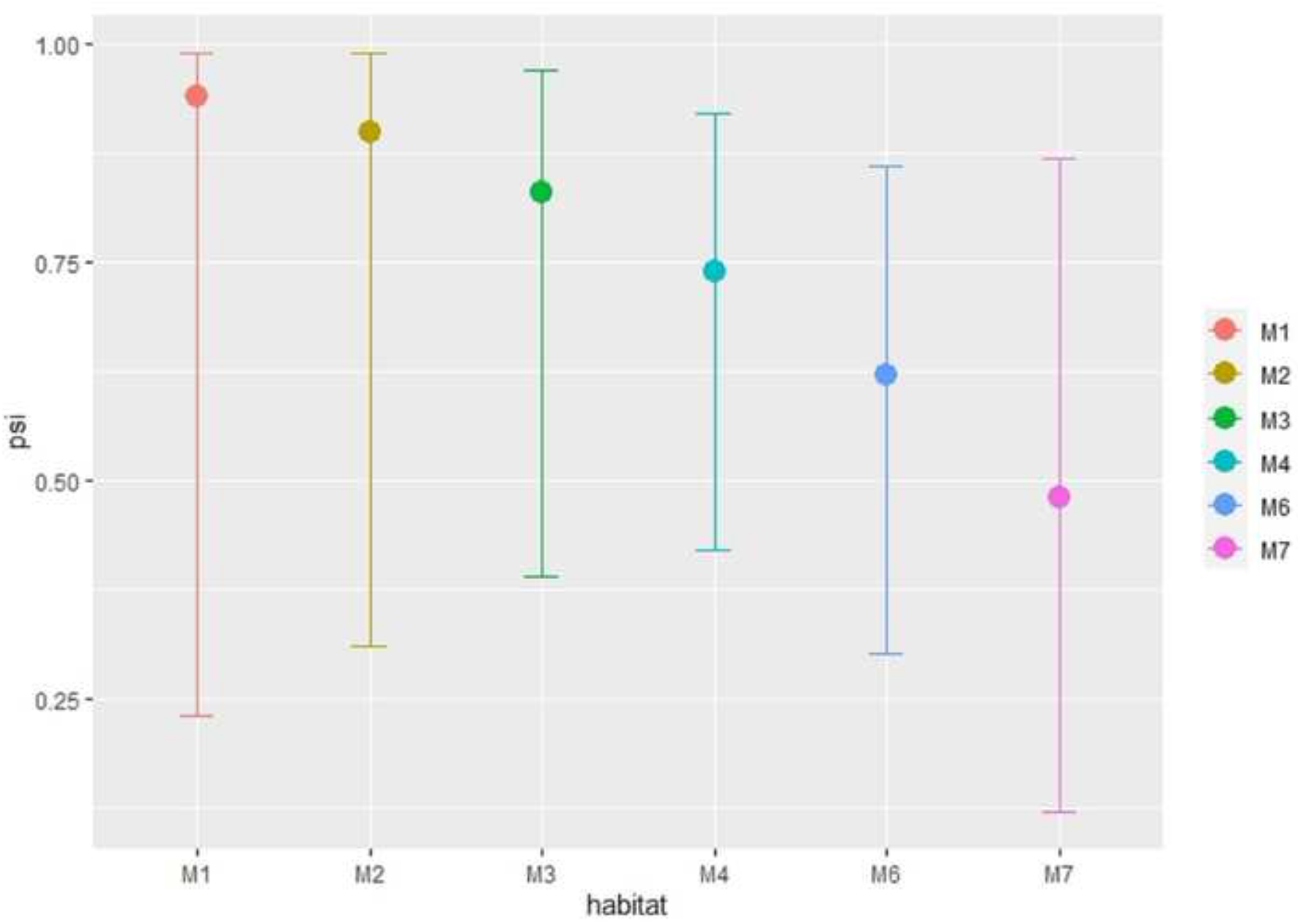
Occupancy 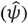 estimates of *Alsophis rufiventris* per habitat based on transect surveys conducted in 2021 on Saba, Caribbean Netherlands. M1 (*Heliconia-Charianthus* Mountains), M2 (*Philodendron-Marcgravia* Mountains), M3 (*Philodendron-Inga* Mountains), M4 (*Swietenia* Mountains), M6 (*Coccoloba-Inga* Mountains), and M7 (*Aristida-Bothriochloa* Mountains).

**Figure 4.**
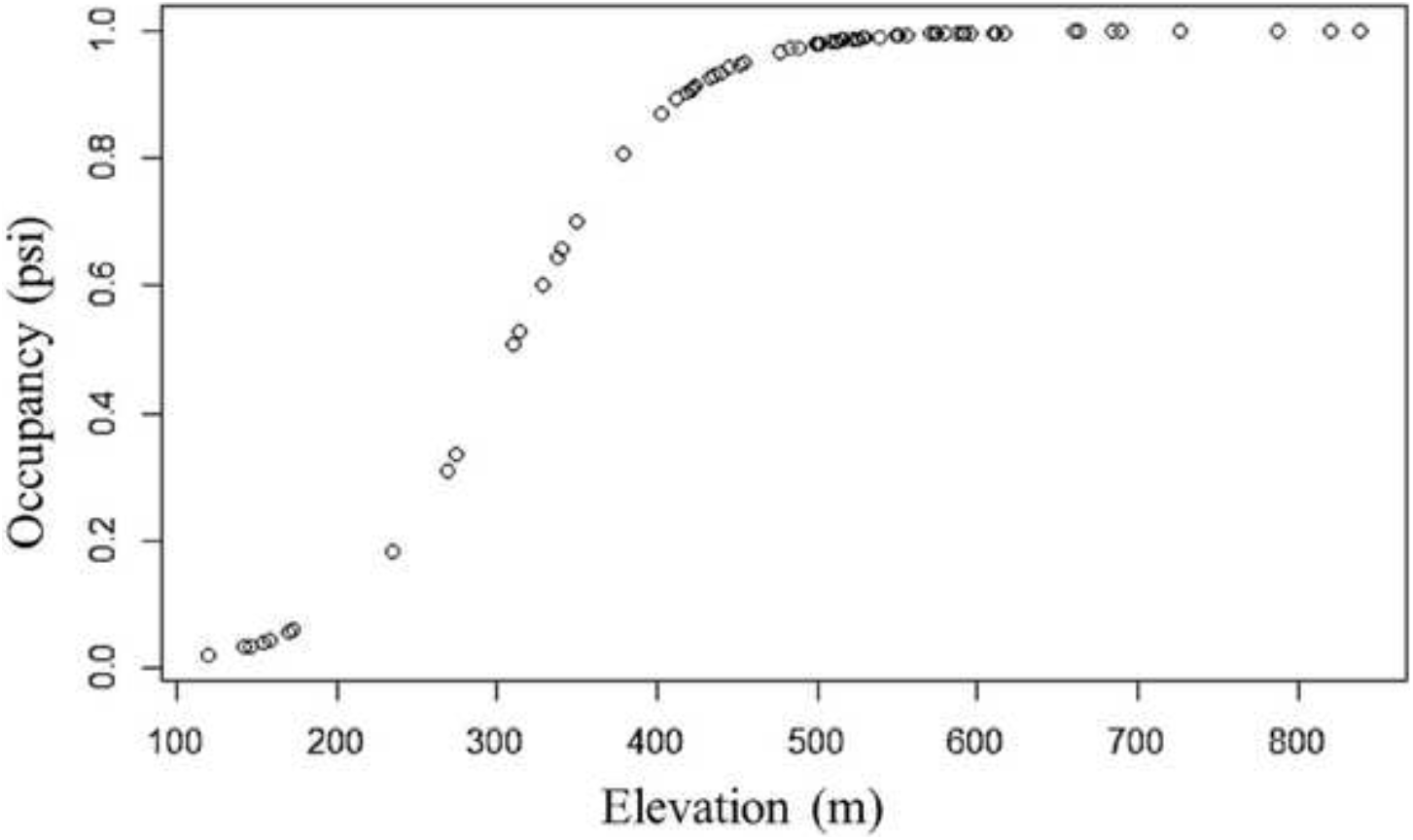
Occupancy 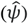 estimates of *Alsophis rufiventris* in relation to elevation (m) based on transect surveys conducted in 2021 on Saba, Caribbean Netherlands.

**Figure 5.**
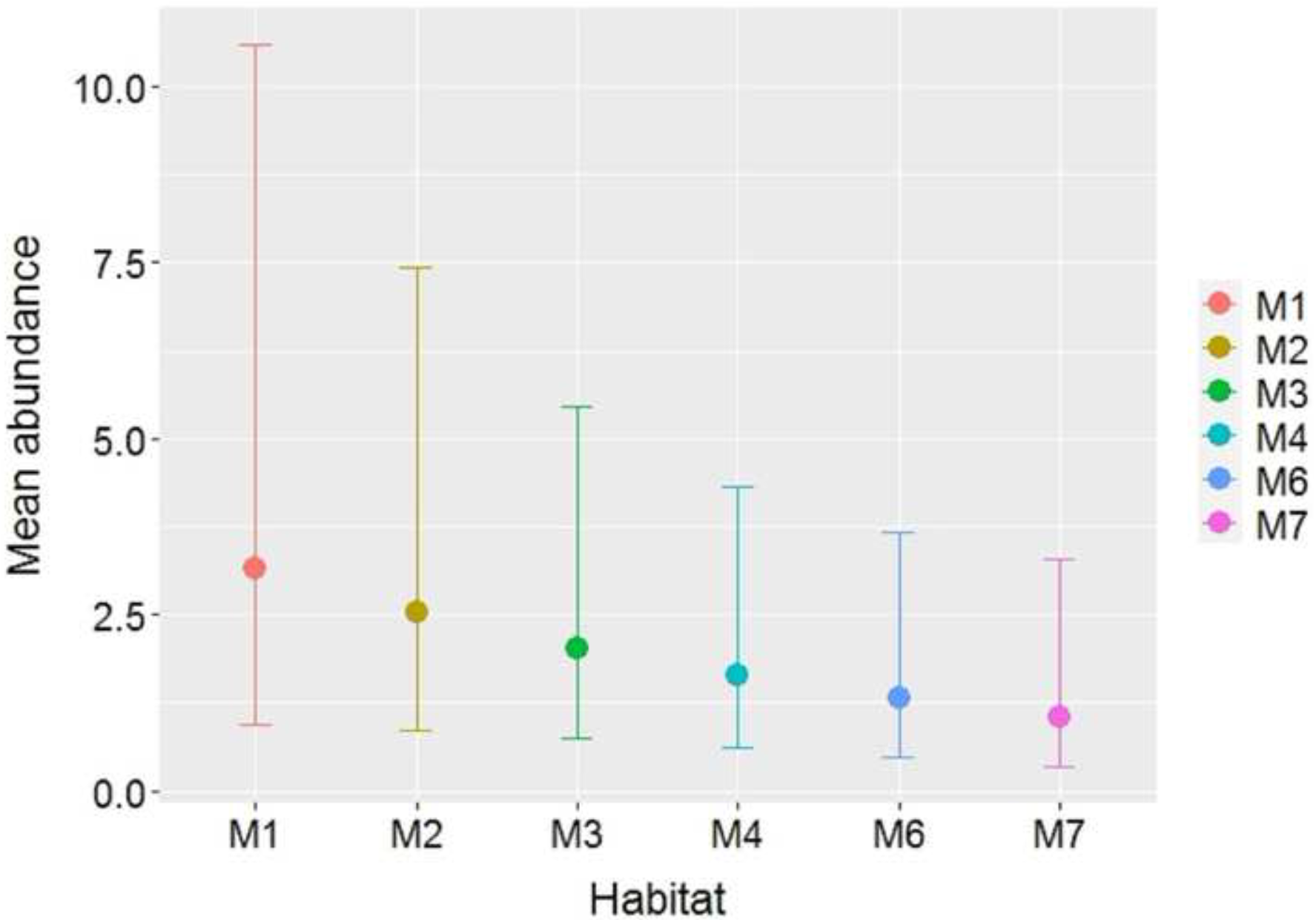
Lambda 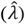 estimates of *Alsophis rufiventris* per habitat based on transect surveys conducted in 2021 on Saba, Caribbean Netherlands. M1 (*Heliconia-Charianthus* Mountains), M2 (*Philodendron-Marcgravia* Mountains), M3 (*Philodendron-Inga* Mountains), M4 (*Swietenia* Mountains), M6 (*Coccoloba-Inga* Mountains), and M7 (*Aristida-Bothriochloa* Mountains).

**Figure 6.**
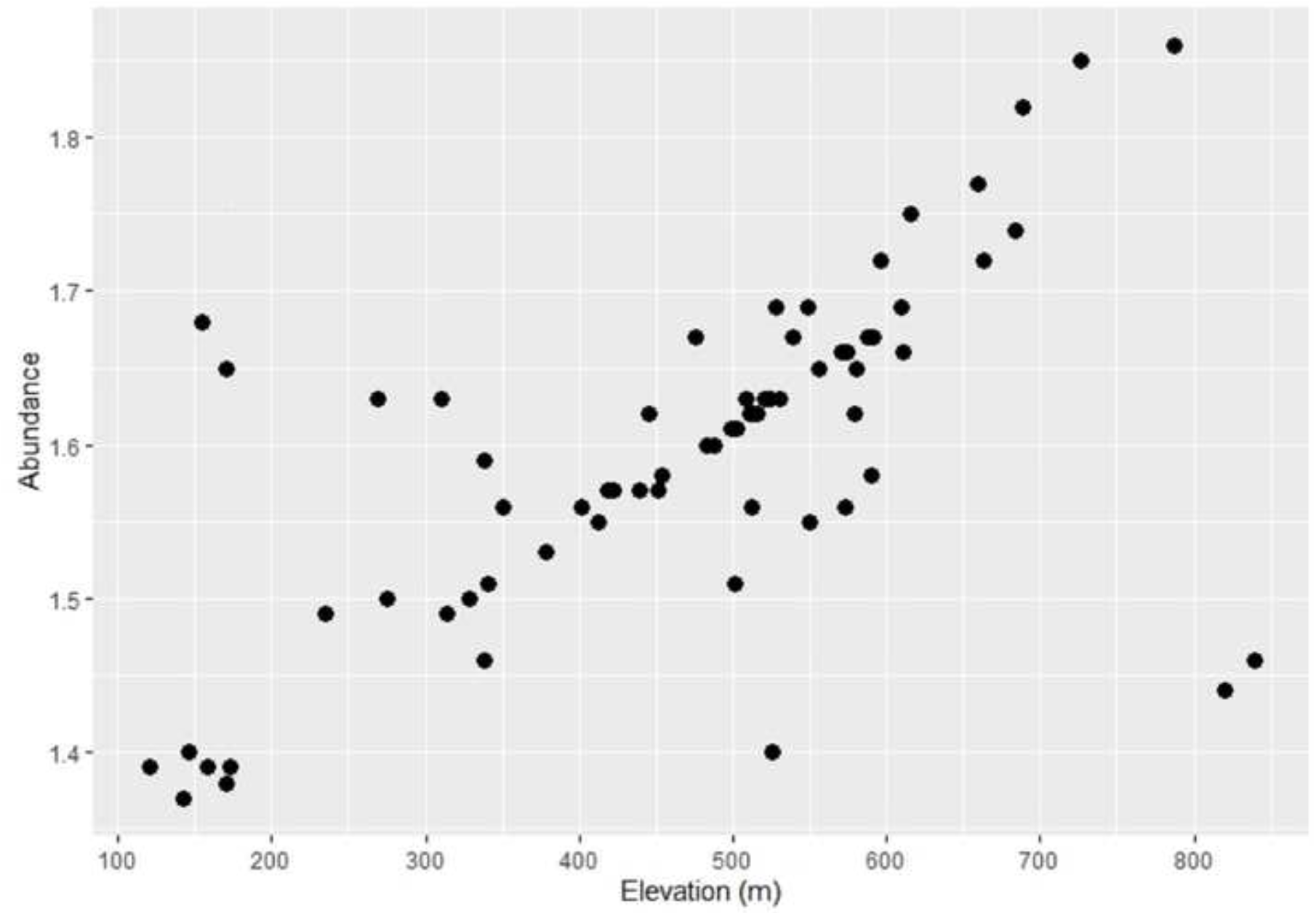
Lambda 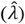 estimates of *Alsophis rufiventris* in relation to elevation (m) based on transect surveys conducted in 2021 on Saba, Caribbean Netherlands.

Using the uniform model with cosine adjustment (Cramer von Mises test: *P* = 0.22), Distance analysis of transect data from Saba revealed 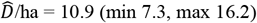, or 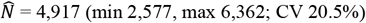 across the entire survey region (438.6 ha). Using the hazard rate model with no adjustment (Cramer von Mises test: *P* = 0.72), Distance analysis of transect data from St. Eustatius in 2018 revealed 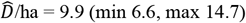, or 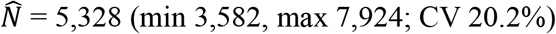 across the entire survey region (540 ha). Distance analysis of transect data from St. Eustatius in 2019 revealed 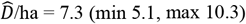, or 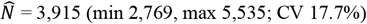.

## DISCUSSION

Due to their secretive nature and cryptic behavior, snakes are among the most difficult vertebrates to study in their native habitats (Durso et al., 2011). On Guam, for example, 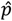 of the brown tree snake (*Boiga irregularis* Bechstein, 1802) is just 0.07 (Christy et al., 2010). We present the first quantitative assessment of *A. rufiventris* on Saba, one of just two islands where this species remains (Saba and St. Eustatius represent 10.9% of the species’ original range; Sajdak and Henderson, 1991). Our results suggest that racers are present in varying densities across the habitats surveyed, and are also present in other habitats on Saba (LM, pers. obs.; K. Wulf, J. Johnson and M. Terpstra, pers. comm.). Although pre-hurricane data are lacking from Saba, density estimates of *Alsophis antiguae* (Parker, 1933) from Great Bird Island (20.2/ha; Daltry et al., 2017) and *Borikenophis portoricensis* (Reinhardt and Lütken, 1862) on Guana Island, British Virgin Islands (19/ha; Hileman et al., 2017) may be representative of a ‘healthy’ racer population on islands such as Saba and St. Eustatius (Hileman et al. even believe their abundance estimate may be underestimated due to the small sample size). In an earlier study, Rodda et al. (2001) surveyed *B. portoricensis* on Guana, where they estimated a density of 50/ha in early successional (*Leucaena leucocephala*) forest habitat, and even suggested this was an underestimate. However, surveys of the St. Lucia Racer (*Erythrolamprus ornatus* Garman, 1887) on Maria Major Island resulted in an estimate of just 8.6/ha (Ross and Williams, 2012), thus racer densities are likely to be species- and habitat-specific. Madden et al. (2021) provided racer density estimates of *A. rufiventris* on St. Eustatius of 9.2/ha (pre-hurricane) and 4.6/ha to 5.0/ha (post-hurricane), which differ from our current estimates. Moreover, Henderson (1992) argues that mongoose-free islands with high human population densities (e.g., Guadeloupe) support higher racer densities compared to those with lower human densities, where in most cases mongoose predation has led to extirpation of the species (e.g., Nevis and Antigua). This is supported by Hileman et al. (2017) who note the abundance of racers on mongoose-free islands.

Our 2021 raw encounter rate is higher than post-hurricane estimates from St. Eustatius but lower than the pre-hurricane estimate from 2011 (Madden et al., 2021), as well as Daltry et al.’s (1997) estimate from Saba (which was based on just a few days’ of fieldwork in March and may not be comparable). Similarly, the higher density estimate from Saba compared to St. Eustatius may be attributed to the fact that surveys on Saba were conducted four years post-hurricane, allowing for recovery of the population; nevertheless, our current density estimate is likely lower than pre-hurricane densities. Unfortunately no data exist to support this speculation and we are cognizant of the limited sample sizes from Saba and St. Eustatius, which limited our ability to produce density and abundance estimates for the different habitats and produced results with levels of precision that are only just considered acceptable (Conquest, 1983). However, Daltry et al. (2017) state that *A. antiguae* can reach high densities, even on small islands, and historical anecdotal observations from Saba confirm this. We note that vegetation types on Saba differ from those on St. Eustatius, and that the understory on Saba is denser at higher elevations (LM, pers. obs.). De Freitas et al. (2016) confirm this by comparing vegetation types on Saba, showing that average vegetation cover is highest (94 %) on the top of Mount Scenery (M1-*Heliconia-Charianthus* Mountains) due to a dense shrub and herb layer while herbs are usually scarce in areas at lower elevations (M4 - *Swietenia* Mountains).

Our results from Saba are consistent with those found by Madden et al. (2021) on St. Eustatius, which suggest that sites are more likely to be occupied at higher elevations (especially >400m). Similarly, mean abundance is predicted to increase in line with elevation, however this drops sharply >800m. Many reptile species have narrowly distributed elevational ranges (Chettri et al., 2010), primarily due to the physiological response of reptiles to temperature (Fu et al., 2007). It may be that the peak of Mount Scenery on Saba, which often ‘snags’ clouds (Powell 2006), is less favorable to racers than habitats at lower elevations, which is supported by Sajdak and Henderson (1991) who did not find racers above 830m. We did not survey any habitats below 100m and as such are unable to comment on the density, occupancy or abundance of racers in these areas, however published and anecdotal observations exist of snakes in urban areas (often killed by road vehicles) and in coastal areas (e.g., Sajdak and Henderson, 1991; Terpstra et al., 2015; LM pers. obs.).

Besides natural disasters, invasive species are cited as a major threat to racers in the genus (Daltry et al., 1997; Powell, 2006). Although Saba (and St. Eustatius) are mongoose-free, both islands have been invaded by black rats and house mice (*Mus musculus* Linnaeus, 1758; Madden et al., 2019, 2020), as well as free-roaming cats and dogs (Rojer 1997). While abundance and distribution data of invasive rodents are lacking from Saba, the impact of cats on the island’s globally significant population of Red-billed Tropicbirds (*Phaethon aethereus* Linnaeus, 1758) is well documented (Debrot et al., 2004; Terpstra et al., 2015; Boeken, 2016). Debrot et al. (2014) estimated 285 cats/km^2^ at one coastal colony based on scat density (1.1% of which contained racers), after which a localized cat removal program was initiated to conserve Tropicbird chicks. No racers were found in the euthanized cats’ stomach contents (*N* = 17), but live racers were documented at the nest colony via camera traps; black rats were also detected in the same area (Terpstra et al., 2015). We are especially concerned about the impacts of free-roaming domestic cats on Saba, where four incidents of racer predation were documented (Fig. 7; LM and B. Noort, pers. obs.) within two weeks.

**Figure 7.**
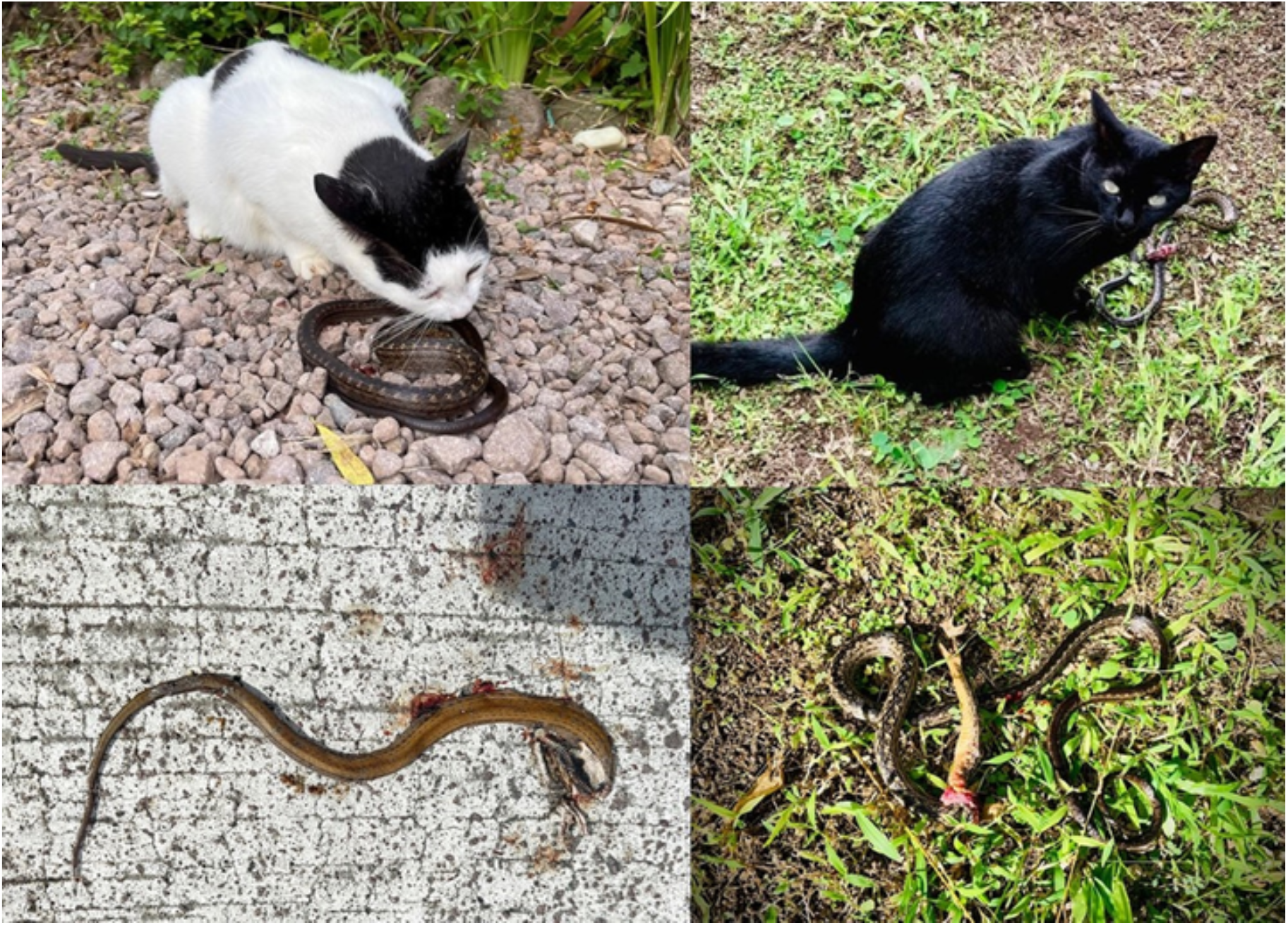
Cat predation of *Alsophis rufiventris* on Saba, Caribbean Netherlands (photos courtesy of Bart Noort).

Despite its small size, Saba supports a number of (regionally) endemic reptile species, including the recently described *Iguana melanoderma* (Breuil et al., 2020), *A. rufiventris* and the endemic *Anolis sabanus* (Garman, 1887), whose population was previously estimated at around seven million individuals (Staats and Schall, 1996). This anole is widely distributed in almost all habitats across the island, described as abundant from sea level to the top of Mount Scenery, and rare only in very dry areas (Lazell, 1972; Rojer, 1997). Anoles and frogs (*Eleutherodactylus johnstonei* Barbour, 1914), which are abundant on Saba and St. Eustatius and whose ranges overlap extensively, form an important food source for the racer (Henderson and Sajdak, 1996). Quantitative data on the impacts of hurricanes on these species are lacking from Saba and St. Eustatius, but studies on other *Anolis* species in the Neotropics have demonstrated that hurricanes can drive natural selection (Donihue et al., 2018, 2020; Dufour et al., 2019). Moreover, the limited landmasses of both islands combined with ongoing habitat destruction/loss and increased hurricanes forces reptile populations into increasingly smaller patches of suitable habitat (Powell 2004). We therefore stress the need to continue monitoring in order to gain a better understanding of temporal trends in *A. rufiventris* populations on both islands, especially considering the ongoing threat of invasive species and the very real risk of another destructive hurricane in the near future (Elsner, 2006).

## ACKNOWLEDGEMENTS

We are grateful to the Saba Conservation Foundation for allowing us to conduct racer surveys on Saba in 2021. We greatly appreciate fieldwork assistance by Tom Wijers and Jasper Bleijenberg. We also thank Kevin Verdel and James Johnson who provided useful practical advice on conducting fieldwork.

## REFERENCES

Alves R.R.N., Vieira K.S., Santana G.G., Vieira W.L.S., Almeida W.O., Souto W.M.S., Montenegro P.F.G.P., Pezzuti J.C.B. 2012. A review on human attitudes towards reptiles in Brazil. Environmental Monitoring and Assessment 184:6877–6901.

Atwood T.B., Valentine S.A., Hammill E., McCauley D.J., Madin E.M., Beard K.H. Pearse W.D. 2020. Herbivores at the highest risk of extinction among mammals, birds, and reptiles. Science advances 6:p.eabb8458.

Barbour T. 1914. A contribution to the Zoogeography of the West Indies, with special reference to amphibians and reptiles. Memoirs of the Museum of Comparative Zoology 44:209–359.

Bechstein J. M. 1802. Herrn de Lacépède’s Naturgeschichte der Amphibien oder der eyerlegenden vierfüssigen Thiere und der Schlangen. Eine Fortsetzung von Buffon’s Naturgeschichte aus dem Französischen übersetzt und mit Anmerkungen und Zusätzen versehen. Weimar: Industrie Comptoir.

Bender M.A., Knutson T.R., Tuleya R.E. et al. 2010. Modeled Impact of Anthropogenic Warning on the Frequency of Intense Atlantic Hurricanes. Science 327:454–458.

Boeken M. 2016. Breeding success of Red-billed Tropicbirds Phaethon aethereus on the Caribbean island of Saba. Ardea 104:263–271.

Breuil M., Schikorski D., Vuillaume B., Krauss U., Morton M.N., Corry E., Bech N., Jelić M., Grandjean F. 2020. Painted black: Iguana melanoderma (Reptilia, Squamata, Iguanidae) a new melanistic endemic species from Saba and Montserrat islands (Lesser Antilles). ZooKeys 926:95.

Buckland S.T., Anderson D.R., Burnham K.P., Laake J.L., Borchers D.L., Thomas L. 2001. Introduction to distance sampling: estimating abundance of biological populations. Oxford University Press, Oxford, UK.

Buckland S.T., Rexstad E.A., Marques T.A., Oedekoven, C.S. 2015. Distance sampling: methods and applications (Vol. 431). New York, NY, USA: Springer.

Catford J.A., Vesk P.A., Richardson D.M., Pyšek P. 2012. Quantifying levels of biological invasion: towards the objective classification of invaded and invasible ecosystems. Global Change Biology 18:44–62. https://doi.org/10.1111/j.1365-2486.2011.02549.x

Central Bureau of Statistics. 2021. Caribbean Netherlands; population; sex, age, marital status. https://www.cbs.nl/en-gb/figures/detail/83698ENG?dl=FB7D. Accessed 21 October 2021.

Chettri B., Bhupathy S., Acharya B.K. 2010. Distribution pattern of reptiles along an eastern Himalayan elevation gradient, India. Acta Oecologica 36:16–22.

Christy M.T., Yackel Adams A.A., Rodda G.H., Savidge J.A., Tyrrell C.L. 2010. Modelling detection probabilities to evaluate management and control tools for an invasive species. Journal of Applied Ecology 47:106–113.

Conquest L.L. 1983. Assessing the statistical effectiveness of ecological experiments: utility of the coefficient of variation. International Journal of Environmental Studies 20:209–221.

Daltry J.C., Day M., Ford N. 1997. Red-bellied Racer conservation project. Conservation status of the Red-bellied Racer. Fauna & Flora International, unpublished report.

Daltry J.C., Lindsay K., Lawrence S.N., Morton M.N., Otto A,. Thibou A. 2017. Successful reintroduction of the Critically Endangered Antiguan racer Alsophis antiguae to offshore islands in Antigua, West Indies. International Zoo Yearbook 51:97–106.

Debrot A.O., Ruijter M., Endarwin W., van Hooft P., Wulf K. 2014. Predation threats to the Red-billed Tropicbird breeding colony of Saba: focus on cats (No. C011/14). IMARES, unpublished report.

de Freitas J.A., Rojer A.C., Nijhof B.S.J., Debrot A.O. 2016. A landscape ecological vegetation map of Saba (Lesser Antilles). Wageningen, IMARES Wageningen UR (University & Research Centre), IMARES, unpublished report C195/15. 48 pp.; 7 tab.

Donihue C.M., Herrel A., Fabre A.-C., Kamath A., Geneva A.J., Schoener T.W., Kolbe J.J., Losos J.B. 2018. Hurricane-induced selection on the morphology of an island lizard. Nature 560:88–91. https://doi.org/10.1038/s41586-018-0352-3.

Donihue C.M., Kowaleski A.M., Losos J.B., Algar A.C., Baeckens S., Buchkowski R.W., Fabre A.-C., Frank H.K., Geneva A.J., Reynolds R.G., Stroud J.T., Velasco J.A., Kolbe J.J., Mahler D.L., Herrel, A. 2020. Hurricane effects on Neotropical lizards span geographic and phylogenetic scales. Proceedings of the National Academy of Sciences of the United States of America, 117:10429–10434. https://doi.org/10.1073/pnas.2000801117.

Dufour C.M., Donihue C.M., Losos J.B., Herrel A. 2019. Parallel increases in grip strength in two species of Anolis lizards after a major hurricane on Dominica. Journal of Zoology 309:77–83. https://doi.org/10.1111/jzo.12685.

Duméril A., Bibron G., Duméril A. 1854. Alsophis rufiventris. Published in: Garman, S. 1887. On West Indian reptiles in the Museum of Comparative Zoology at Cambridge, Mass. Proceedings of the American Philosophical Society 24:278–286.

Durso A.M., Willson J.D., Winne C.T. 2011. Needles in haystacks: estimating detection probability and occupancy of rare and cryptic snakes. Biological Conservation 144:1508–1515.

Efron B., Tibshirani R.J. 1993. An introduction to the bootstrap. Monographs on Statistics and Applied Probability 57:1–436.

Elsner J.B. 2006. Evidence in support of the climate change–Atlantic hurricane hypothesis. Geophysical Research Letters 33.

Fauth J.E., Crother B.I., Slowinski J.B. 1989. Elevational patterns of species richness, evenness, and abundance of the Costa Rican leaf-litter herpetofauna. Biotropica:178–185.

Fu C., Wang J., Pu Z., Zhang S., Chen H., Zhao B., Chen J., Wu J. 2007. Elevational gradients of diversity for lizards and snakes in the Hengduan Mountains, China. Biodiversity Conservation 16:707–726.

Garman S. 1887. On West Indian reptiles in the Museum of Comparative Zoology at Cambridge, Mass. Proceedings of the American Philosophical Society 24:278–286.

Geoffroy Saint-Hilaire, É. 1818. Herpestes javanicus. In Species 2000 & ITIS Catalogue of Life: 2019, Catalogue of Life.

Gibbons J.W., Scott D.E., Ryan T.J., Buhlmann K.A., Tuberville T.D., Metts B.S., Greene J.L., Mills T., Leiden Y., Poppy S., Winne C.T. 2000. The global decline of reptiles, déjà vu amphibians. BioScience 50:653–666.

Henderson R.W. 1992. Consequences of predator introductions and habitat destruction on amphibians and reptiles in the post-Columbus West Indies. Caribbean Journal of Science 28:1–10.

Henderson R.W., Sajdak R.A. 1996. Diets of West Indian racers (Colubridae: Alsophis): composition and biogeographic implications. Contributions to West Indian Herpetology: A Tribute to Albert Schwartz:327–338.

Henderson R.W. 2004. Lesser Antillean snake faunas: Distribution, ecology, and conservation concerns. Oryx 38:311–320.

Herrera-Montes A., Brokaw N. 2010. Conservation value of tropical secondary forest: A herpetofaunal perspective. Biological Conservation 143:1414–1422.

Hileman E.T., Powell R., Perry G., Mougey K., Thomas R., Henderson R.W. 2017. Demography of the Puerto Rican Racer, Borikenophis portoricensis (Squamata: Dipsadidae), on Guana Island, British Virgin Islands. Journal of Herpetology 51:454–460.

Lazell J.D., Jr. 1972. The anoles (Sauria, Iguanidae) of the Lesser Antilles. Bulletin of the Museum of Comparative Zoology 143:1–115.

Linnaeus C. 1758. Systema naturae per regna tria naturae, secundum classes, ordines, genera, species, cum characteribus, differential, synonymis, locis, Tomus I. Editio decima, reformata.

Lugo A.E. 2000. Effects and outcomes of Caribbean hurricanes in a climate change scenario. Science of the Total Environment 262:243–251.

MacKenzie D.I., Nichols J.D., Royle J.A., Pollock K.H., Bailey L., Hines J.E. 2017 Occupancy estimation and modeling: inferring patterns and dynamics of species occurrence. Elsevier, Amsterdam.

Madden H., Van Andel T., Miller J., Stech M., Verdel K., Eggermont E. 2019. Vegetation associations and relative abundance of rodents on St. Eustatius, Caribbean Netherlands. Global Ecology and Conservation 20:p.e00743.

Madden H., Eggermont E., Verdel K. 2020. Micro-and Macrohabitat Preferences of Invasive Rodents on St. Eustatius, Caribbean Netherlands. Caribbean Journal of Science 50:202–211.

Madden H. 2020. Nocturnal observation of Alsophis rufiventris. Natural History Notes. Herpetological Review 51.

Madden H., Fernández D.S., Tremblay R.L., Verdel K., Kaboord B. 2021. Find me if you can: Pre- and post-hurricane densities of the Red-bellied Racer (Alsophis rufiventris) on St. Eustatius, and a review of the genus in the Caribbean. bioRxiv. https://doi.org/10.1101/2021.07.05.451169.

Maley A. 2004. Species profile: Red-bellied Racer (Alsophis rufiventris). Iguana 11:147.

Maley A.J., Savit A.Z., Heinz H.M., Powell R., Henderson R.W. 2006. Alsophis rufiventris. Catalogue of American Amphibians and Reptiles (CAAR).

Marques T.A., Thomas L., Fancy S.G., Buckland S.T. 2007. Improving estimates of bird density using multiple-covariate distance sampling. The Auk 124(4):1229–1243.

Mayani-Parás F., Botello F., Castañeda S., Sánchez-Cordero V. 2019. Impact of habitat loss and mining on the distribution of endemic species of amphibians and reptiles in Mexico. Diversity 11:210.

McDonald T., Carlisle J., McDonald A. 2019. Rdistance: Distance-Sampling Analyses for Density and Abundance Estimation. R package version 2.1.3. https://CRAN.R-project.org/package=Rdistance.

Meredith M. 2020. wiqid: Quick and Dirty Estimates for Wildlife Populations. R package version 0.3.0. Available from https://CRAN.R-project.org/package=wiqid.

Mittermeier R.A., Robles G.R., Hoffman M., Pilgrim J., Brooks T., Mittermeier C.G., Lamoreux J., da Fonseca G.A.B. 2004. Hotspots revisited: Earth’s biologically richest and most threatened terrestrial ecoregions. CEMEX, Mexico DF.

Moser D., Lenzner B., Weigelt P., Dawson W., Kreft H., Pergl J., Pyšek P., van Kleunen M., Winter M., Capinha C., Cassey P. 2018. Remoteness promotes biological invasions on islands worldwide. Proceedings of the National Academy of Sciences 115:9270–9275.

Parker H.W. 1933. Some amphibians and reptiles from the Lesser Antilles. Annals of the Magazine of Natural History 11:151–158.

Powell R. 2004. Conservation of iguanas (Iguana delicatissima and I. iguana) in the Lesser Antilles. Iguana 11:238–246.

Powell R., Henderson R.W. 2005. Conservation status of Lesser Antillean reptiles. Iguana 12:62–77.

Powell R. 2006. Conservation of the herpetofauna on the Dutch Windward Islands: St. Eustatius, Saba, and St. Maarten. Applied Herpetology 3:293–306.

Questel K. 2021. Alsophis rijgersmaei Cope, 1869 (Squamata: Dipsadidae) sur l’île de Saint-Barthélemy. Agence Territoriale de l’Environnement St. Barthélemy.

Reinhardt J., Lütken C.F. 1862. Bidrag tii det vestindiske Öriges og navnligen tii de dansk-vestindiske Oers Herpetologie. Videnskabelige meddelelser fra den Naturhistoriske forening i Kjöbenhavn:153–291.

Ricklefs R.E., Lovette I.J. 1999. The roles of island area per se and habitat diversity in the species–area relationships of four Lesser Antillean faunal groups. Journal of Animal Ecology 68:1142–1160.

Rodda G.H., Perry G., Rondaeu R.J., Lazell J. 2001. The densest terrestrial vertebrate. Journal of Tropical Ecology 17:331–338.

Rojer A. 1997. Biological inventory of Saba. Curaçao, Netherlands Antilles: Carmabi Foundation.

Ross T., Williams R.J. 2012. Population assessment and conservation status of the Saint Lucia racer, Liophis ornatus. November - December 2011. Unpubl. report to Durrell Wildlife Conservation Trust, Saint Lucia Forestry Department, Saint Lucia National Trust and Fauna & Flora International.

Sajdak R.A., Henderson R.W. 1991. Status of West Indian racers in the Lesser Antilles. Oryx 25:33–38.

Staats C.M., Schall J.J. 1996. The Saban Anole: the story of a unique lizard on a small Caribbean island. Saba Conservation Foundation, Saba, Netherlands Antilles.

Stoffers A.L. 1956. The vegetation of the Netherlands Antilles (No. 135). Kemink.

Terpstra M., Van der Woude E., Wulf K., van Rijn J., Debrot A.O., 2015. Monitoring the effect of cat removal on reproductive success in Red-billed Tropicbird colonies on Saba, 2013-2014: first season of results (No. C103/15). IMARES.

Urban M.C. 2015. Accelerating extinction risk from climate change. Science 348:571–573.

Veenenbos J.S. 1955. A soil and land capability survey of St. Maarten, St. Eustatius, and Saba. Publ. Found. Scientific Research Suriname Netherlands Antilles, Utrecht, The Netherlands. 11, 94 pp., ill., map

Webster P.J., Holland G.J., Curry J.A., Chang H.R. 2005. Changes in tropical cyclone number, duration, and intensity in a warming environment. Science 309:1844–1846.

